# Pharmacokinetically optimized anesthesia enables long-term functional ultrasound imaging in the cat visual cortex

**DOI:** 10.64898/2026.08.24.746778

**Authors:** Domonkos Horváth, Klaudia Csikós, Ábel Petik, Attila Balázs Dobos, Daniel Hillier

## Abstract

Functional ultrasound imaging (fUSI) measures cerebral blood-volume responses, so anesthesia protocols developed for BOLD fMRI may not preserve its signal. We screened three fMRI-derived anesthesia regimens for visually evoked fUSI in the cat visual cortex. Isoflurane– ketamine–medetomidine produced the strongest and most consistent responses, but the standard intramuscular medetomidine bolus suppressed the signal in one sensitive cat. Pharmacokinetic modeling guided replacement of this bolus with controlled intravenous dosing, restoring the visual response while maintaining physiological stability. The same weight-based regimen produced robust responses in the other animals. In a direct within-session test, response strength was equivalent in recordings separated by more than three hours. Across 464 recordings from 51 sessions in three cats, it showed no temporal drift during follow-up extending to 23 months. Visually evoked activation also remained clear relative to a small awake dataset, although equivalence was not established. PK-guided control of medetomidine exposure therefore enables stable, repeated fUSI of the cat visual cortex over hours to years.

## Introduction

Functional ultrasound imaging (fUSI) is an emerging neuroimaging technique that enables high-resolution mapping of brain activity through stimulus-evoked changes in cerebral blood volume (CBV) (Macé et al. 2011; Montaldo et al. 2022). With spatial resolution on the order of 150 µm and imaging rates of up to approximately 10 Hz, fUSI offers substantially higher spatiotemporal resolution than standard functional magnetic resonance imaging (fMRI) in large-animal models. Its portability and compatibility with chronic cranial preparations further make fUSI well suited for repeated measurements in the same animal. Such longitudinal within-animal designs can capture functional changes over timescales ranging from hours to months, reduce variability arising from inter-individual differences, and decrease the number of animals required to study long-term changes in cortical function.

Realizing these advantages requires an experimental preparation that provides stable physiological conditions and reproducible hemodynamic responses across both individual recording sessions and repeated procedures. Awake fUSI has been successfully implemented in large-brained species, including ferrets and nonhuman primates (Bimbard et al. 2018; Blaize et al. 2020). However, awake recordings generally require extensive habituation or behavioral training, and the duration of stable data acquisition can be limited by animal movement and welfare considerations. Repeated anesthetized recordings therefore provide an important complementary strategy, particularly for experiments requiring multi-hour acquisitions, consistent stimulus delivery, gaze control, or sequential sampling of large cortical volumes. For longitudinal fUSI, however, anesthesia must not only maintain an appropriate depth of anesthesia and physiological stability within a session, but must also preserve neurovascular coupling and produce reproducible CBV responses across sessions in the same animal.

Developing such a protocol is not straightforward because anesthetic agents can alter cerebral vascular tone, cerebral perfusion, and neurovascular coupling (Gao et al. 2016; Sumiyoshi, Keeley, and Lu 2019; Bini et al. 2023). Although anesthesia protocols have been established for functional imaging in cats, macaques, and marmosets (Logothetis et al. 2001; Brown et al. 2014; Ortiz-Rios et al. 2024), these protocols have primarily been evaluated using fMRI. Classical blood-oxygen-level-dependent fMRI measures changes related to the oxyhemoglobin-to-deoxyhemoglobin ratio, whereas fUSI measures local changes in CBV (Ogawa et al. 1990; Montaldo et al. 2022). Because the two modalities reflect partially distinct components of the hemodynamic response, an anesthesia protocol that preserves the fMRI BOLD signal cannot be assumed to provide stable or reproducible fUSI responses. A dedicated anesthesia protocol must therefore be evaluated against the specific requirements of longitudinal fUSI.

In this study, we sought to develop an anesthesia protocol that enables stable, reproducible, and longitudinal recording of visually evoked fUSI responses in a chronic cat preparation. We initially screened three published anesthesia methods in a small number of pilot sessions to identify which provided the most robust fUSI signal in our hands. This comparison was not designed as a powered, systematic evaluation of each protocol, but as a pragmatic screening step. As a result of the screening, isoflurane-ketamine-medetomidine anesthesia was found to provide the most reliable fUSI responses among the tested protocols. However, it is well known (Lamont et al. 2001; Pypendop et al. 2011) that medetomidine, especially in combination with inhalational anesthetics such as isoflurane, can induce severe bradycardia (heart rate < 100 bpm) in sensitive animals. In medetomidine-sensitive cats, this cardiovascular effect reduces cerebral perfusion and thereby suppresses the CBV signal underlying fUSI — an outcome we observed in one of our experimental animals.

To overcome the suppressive effects of medetomidine on fUSI signal in sensitive animals, we implemented a pharmacokinetic (PK) model of medetomidine plasma concentration. Medetomidine dosing was then optimized using the fUSI signal as a real-time readout. As a result, we developed a medetomidine-infusion-based isoflurane-ketamine-medetomidine anesthesia protocol that preserves fUSI responses in medetomidine-sensitive cats and generalizes to normal animals.

The optimal anesthesia protocol enables stable fUSI recordings for several hours from the cat visual cortex, repeatable across recording sessions for several months in the same animal.

Finally, as an initial, exploratory characterization, we compared fUSI responses recorded with the optimal anesthesia protocol to those recorded in awake, untrained animals. Because the animals were not trained for head-fixed awake recording, the awake dataset was limited in size, and this comparison was not designed as a powered equivalence study. The observed data are numerically consistent with preserved anesthetized responses, but formal equivalence could not be statistically established due to the limited awake sample size. These results nonetheless provide a preliminary indication of the protocol’s potential for longitudinal fUSI studies in which anesthesia serves as a practical and humane alternative to extensive behavioral training.

## Methods

Before all anesthetized procedures, the cats received antiemetic treatment (maropitant, 1 mg/kg SC, Prevomax, Dechra) in the evening before the anesthetized experiment.

### Animals

Subjects were three adult domestic cats (2 females, 2-4 years). All cats were bred in the animal house of the HUN-REN Research Centre for Natural Sciences. Before the fUSI experiments, a 3D-printed skull chamber was implanted into each cat under general surgical anesthesia and aseptic conditions. 2 weeks after the chamber implantation, under general surgical anesthesia and aseptic conditions, a 30*30 mm craniotomy was drilled leaving the dura mater intact. Then, the craniotomy was filled with a medical-grade, addition-curing silicone earmold material (Biopor® AB, 25 Shore A; Dreve Otoplastik GmbH). The implanted chamber was closed with a 3D-printed cap and screws and sealed with the same silicone until the first fUSI recording 2 weeks after the craniotomy surgery. All procedures were approved by the Animal Care Committee of the HUN-REN Research Centre for Natural Sciences and by the National Food Chain Safety Office of Hungary.

### Isoflurane-ketamine-medetomidine anesthesia

This protocol was adapted from a published anesthesia protocol for fMRI reocrdings in adult cats (Brown et al. 2014). On the experiment day, the cat was premedicated with 0.25 mg/kg midazolam (Dormicum, Egis) and 20 µg/kg medetomidine (Sedator, Dechra) injected intramuscularly. After the loss of reflexes, an intravenous cannula was inserted into one of the cephalic or saphenous veins. Anesthesia was induced by 3.5 mg/kg ketamine (CP-Ketamine, Produlab Pharma) injected intravenously. After intubation, the cat was put in a custom-built head holder that connects to the 3D-printed skull implant and enables pain-free head fixation. Anesthesia was maintained with 1% isoflurane and 0.6 mg/kg/h ketamine infusion during the installation of the functional ultrasound transducer. Isoflurane rate was reduced to 0.5% and ketamine rate was increased to 0.72-0.84 mg/kg/h for fUSI. The first fUSI recording was performed at least 10 minutes after changing the isoflurane and ketamine rates for imaging.

### Alfaxalone-ketamine-medetomidine anesthesia

On the experiment day, the cat was premedicated with 0.25 mg/kg midazolam (Dormicum, Egis) and 20 µg/kg medetomidine (Sedator, Dechra) injected intramuscularly. After the loss of reflexes, an intravenous cannula was inserted into one of the cephalic or saphenous veins. Anesthesia was induced by 2 mg/kg alfaxalone (Alfaxan Multidose, Jurox) injected intravenously. After intubation, the cat was put in a custom-built head holder that connects to the 3D-printed skull implant and enables pain-free head fixation. Anesthesia was maintained with 2-3 mg/kg/h alfaxalone and 0.6 mg/kg/h ketamine infusion during the installation of the functional ultrasound transducer and for fUSI. Medetomidine was administered only as a single premedication dose and was not continued as an infusion.

### Isoflurane-ketamine-fentanyl anesthesia

On the experiment day, the cat was premedicated with 0.25 mg/kg midazolam (Dormicum, Egis) and 3.5 mg/kg ketamine (CP-Ketamine, Produlab Pharma) injected intramuscularly. After the loss of reflexes, an intravenous cannula was inserted into one of the cephalic or saphenous veins. Anesthesia was induced by 1 mg/kg alfaxalone (Alfaxan Multidose, Jurox) and 4 µg/kg fentanyl (Fentanyl Kalceks, As Kalceks) injected intravenously. Alfaxalone was used only for anesthesia induction and not for maintenance. The first recording happened at least 1 hour after alfaxalone injection, thus having little effect on the imaging results, given the short half-life of alfaxalone in cats after intravenous injection (Whittem et al. 2008). After intubation, the cat was put in a custom-built head holder that connects to the 3D-printed skull implant and enables pain-free head fixation. Anesthesia was maintained with 1% isoflurane, 0.6 mg/kg/h ketamine and 2-3 µg/kg/h fentanyl infusion during the installation of the functional ultrasound transducer. Isoflurane rate was reduced to 0.5% and ketamine rate was increased to 0.72-0.84 mg/kg/h for fUSI. The first fUSI recording was performed at least 10 minutes after changing the isoflurane and ketamine rates for imaging.

### Gaze fixation and anesthesia monitoring

To fixate the gaze of the cat on the visual stimulation screen, neuromuscular blockade was induced in all anesthesia protocols with 0.6 mg/kg rocuronium (Rocuronium bromide hameln, Hameln Pharma) injected intravenously and maintained with 0.48-0.8 mg/kg/h rocuronium infusion. Mechanical ventilation was provided in volume-controlled mode (Vt=13 ml/kg, RR=12-20/min) with a veterinary ventilator (R419, RWD). Heart rate, electrocardiogram, blood oxygen saturation, respiratory rate, exhaled carbon-dioxide pressure, non-invasive blood pressure and body temperature were continuously monitored during the anesthetized procedures. Body temperature was maintained 37.5-38.5°C using a thermo-foil blanket and heating pads under and on the cat. The right eye of the animal was treated with a cornea-protective gel and covered with a cotton disc. In the left eye, the nictitating membrane was retracted by phenylephrine, and the pupil was dilated by atropine drops. The eyelids were retracted with an eye speculum, and the cornea was repeatedly moistened with sterile saline. The area centralis was mapped on the center of the screen as described in (Vakkur et al. 1963).

### Awake preparation

For awake fUSI, the untrained cat was put in an open box in front of the visual stimulation screen. The ultrasound transducer was attached to the skull implant, and no head or gaze fixation was applied. The cat was gently held and turned towards the visual stimulation screen during stimulus presentation. The maximum duration of awake recordings was 1 hour.

### Functional ultrasound imaging

All experiments were recorded using a custom-built ultrasound imaging system (Urban et al., 2015). Images were acquired using a linear ultrasound probe (128 elements, 12 MHz) driven by an ultrasound research scanner (Vantage 256, Verasonics) at 1 kHz and processed to yield Doppler images at a frame rate of either 2 or 5 Hz. The ultrasound transducer was positioned above the visual cortex of the cats in a sagittal orientation. The field-of-view covered 16 mm along the cortical surface and 10 mm in depth. The in-plane voxel dimensions were 125 μm (width) × 60 μm (height), and the plane thickness was 400 μm.

### Visual stimulation

Visual stimuli were generated using custom Python software building on the PsychoPy package (Peirce 2007) and displayed on a 55” OLED screen (LG OLED55C41) positioned 42.6 cm in front of the animal’s eyes.

To compare anesthesia methods to awake recordings, as well as for isoflurane-ketamine-medetomidine anesthesia protocol optimization (Figures 1-2, 4), visual stimulation was an alternating full screen checkerboard pattern. The nominal temporal frequency of the alternations was 7 Hz (true display frequency was ∼6.7 Hz due to OLED screen refresh rate), and the spatial frequency of the checkerboard pattern was 0.125 cycles/visual degree. The alternating checkerboard pattern was presented for 4 sec. A 5 sec uniform gray screen before the stimulus and a 10 sec uniform gray screen after the stimulus were presented as a baseline. This stimulation was presented 10 times in each recording, and the temporal averages of the 10 repetitions were used for further analysis.

**Figure 1.**
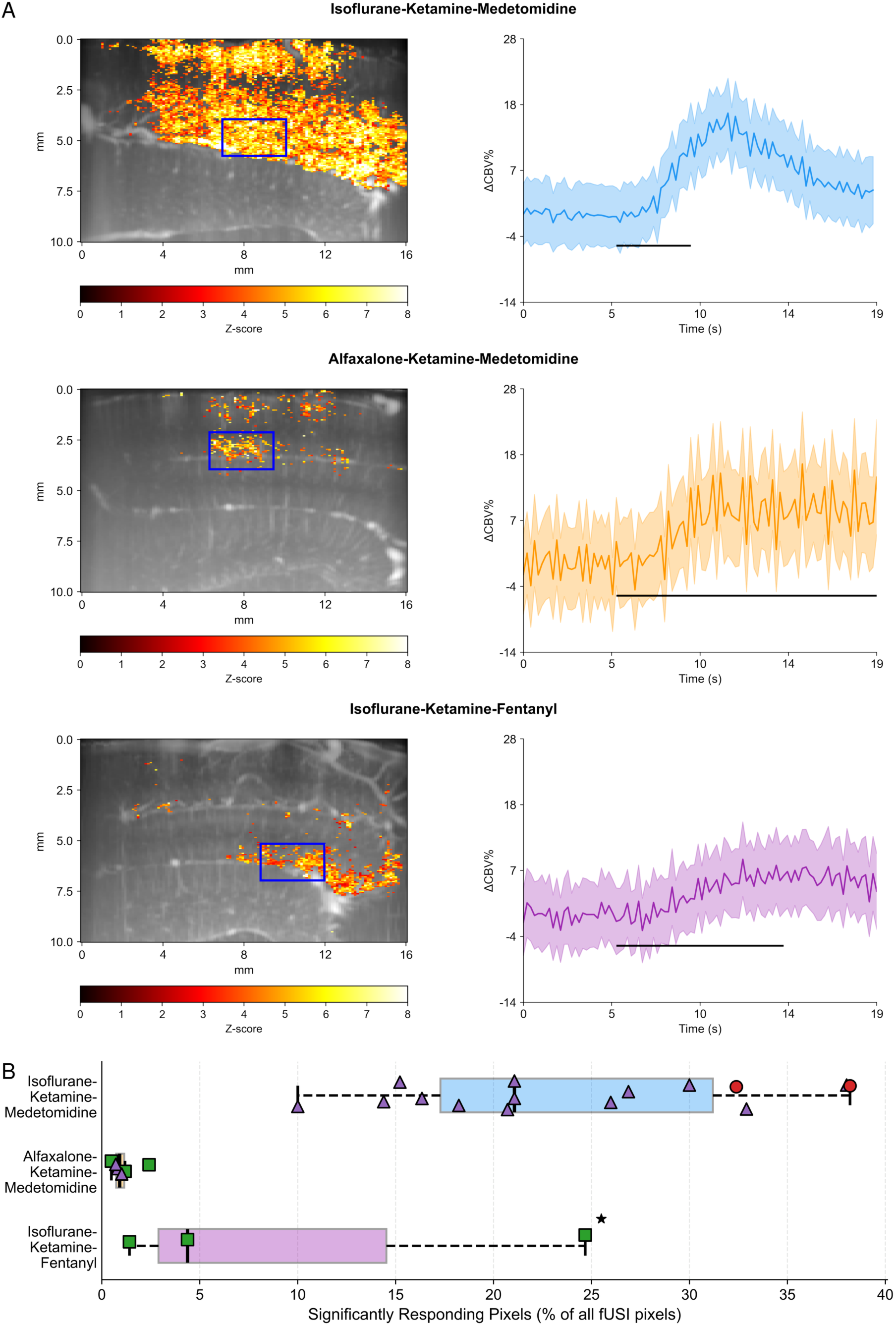
Comparison of visually evoked functional ultrasound imaging responses under different anesthesia protocols in the cat visual cortex. **(A)** Isoflurane-ketamine-medetomidine (top), alfaxalone-ketamine-medetomidine (middle) and isoflurane-ketamine-fentanyl (bottom) anesthesia response maps and average hemodynamic response time courses elicited by a full-screen flickering checkerboard stimulus. Blue rectangles represent the ROIs used to calculate the average CBV response curves in the right column. Horizontal black lines on average CBV response curves represent the visual stimulus duration which was different for each anesthesia protocol. **(B)** Number of significantly responding pixels as percent of all pixels (166*128=21,248) in a fUS image in the different anesthetized conditions. Boxes show median and interquartile range (IQR, 25th–75th percentile); whiskers extend to the most extreme value within 1.5×IQR of the box edges; outliers beyond this range are not displayed. Individual points show single recordings, jittered vertically and coded by experimental animal; n = 13 recordings/12 days isoflurane-ketamine-medetomidine, n = 6 recordings/2 days alfaxalone-ketamine-medetomidine and n = 3 recordings/1 day isoflurane-ketamine-fentanyl, three animals). Asterisk marks the recording where 0.3% isoflurane was used instead of 0.5% for all other recordings.

**Figure 2.**
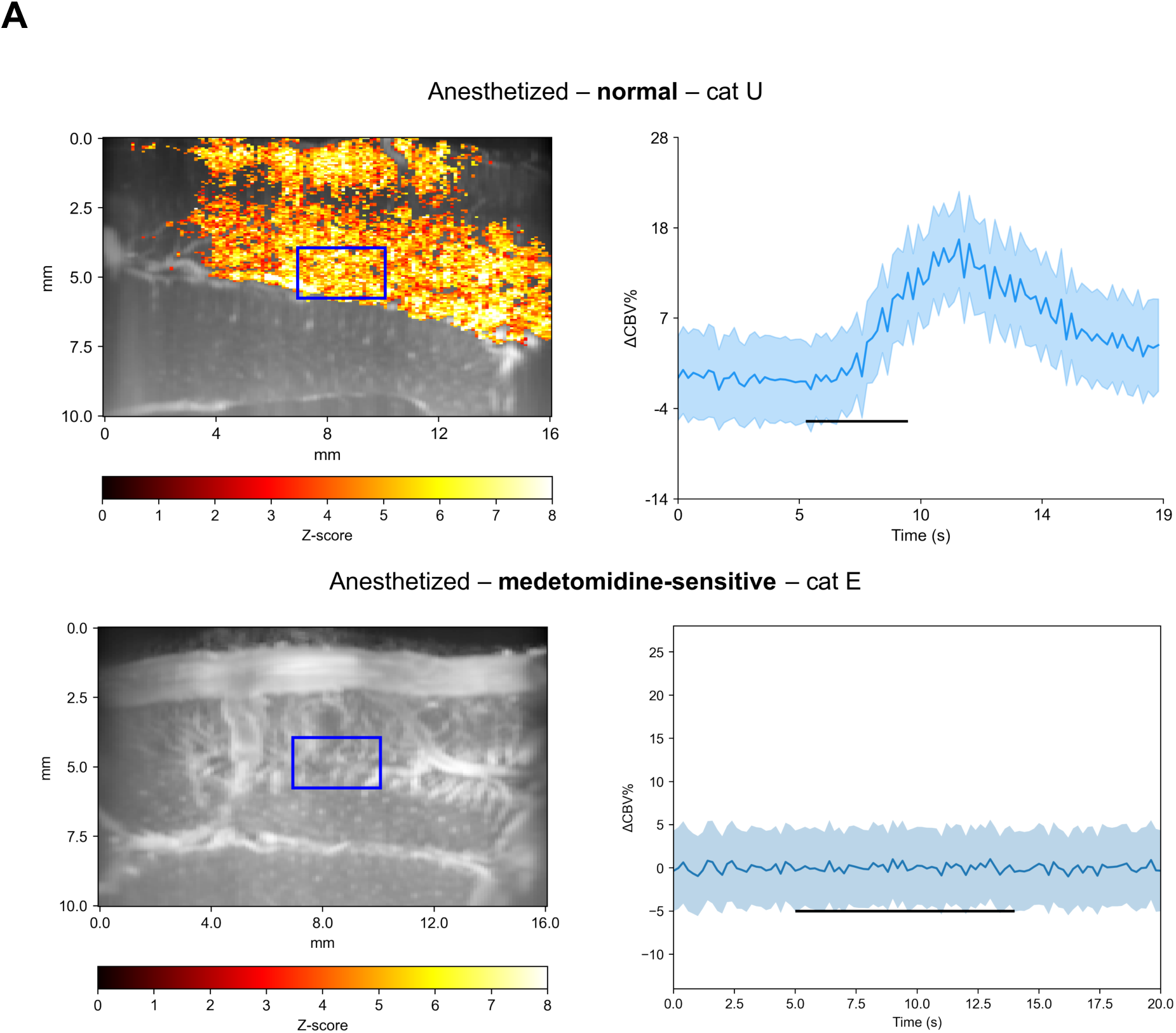
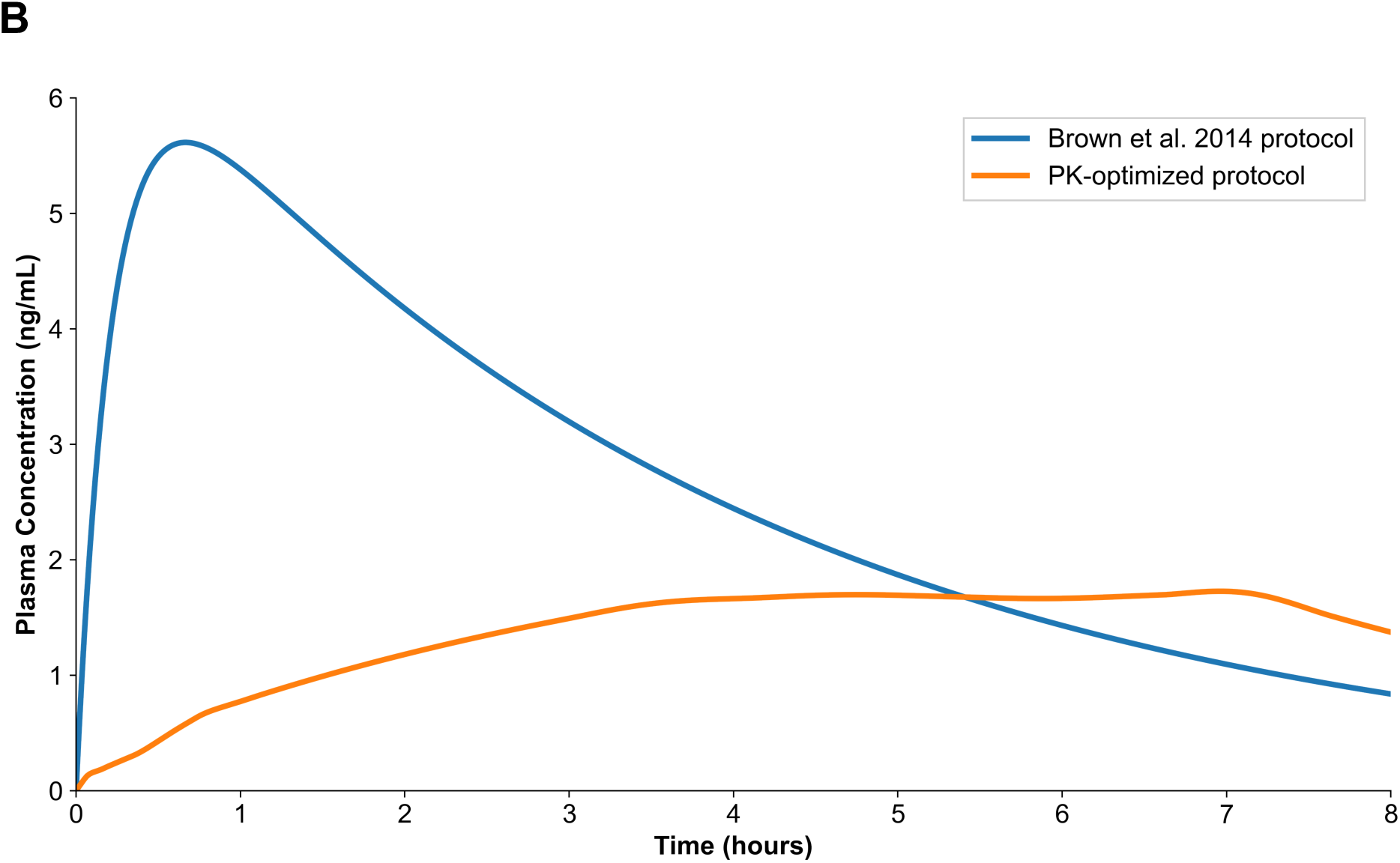
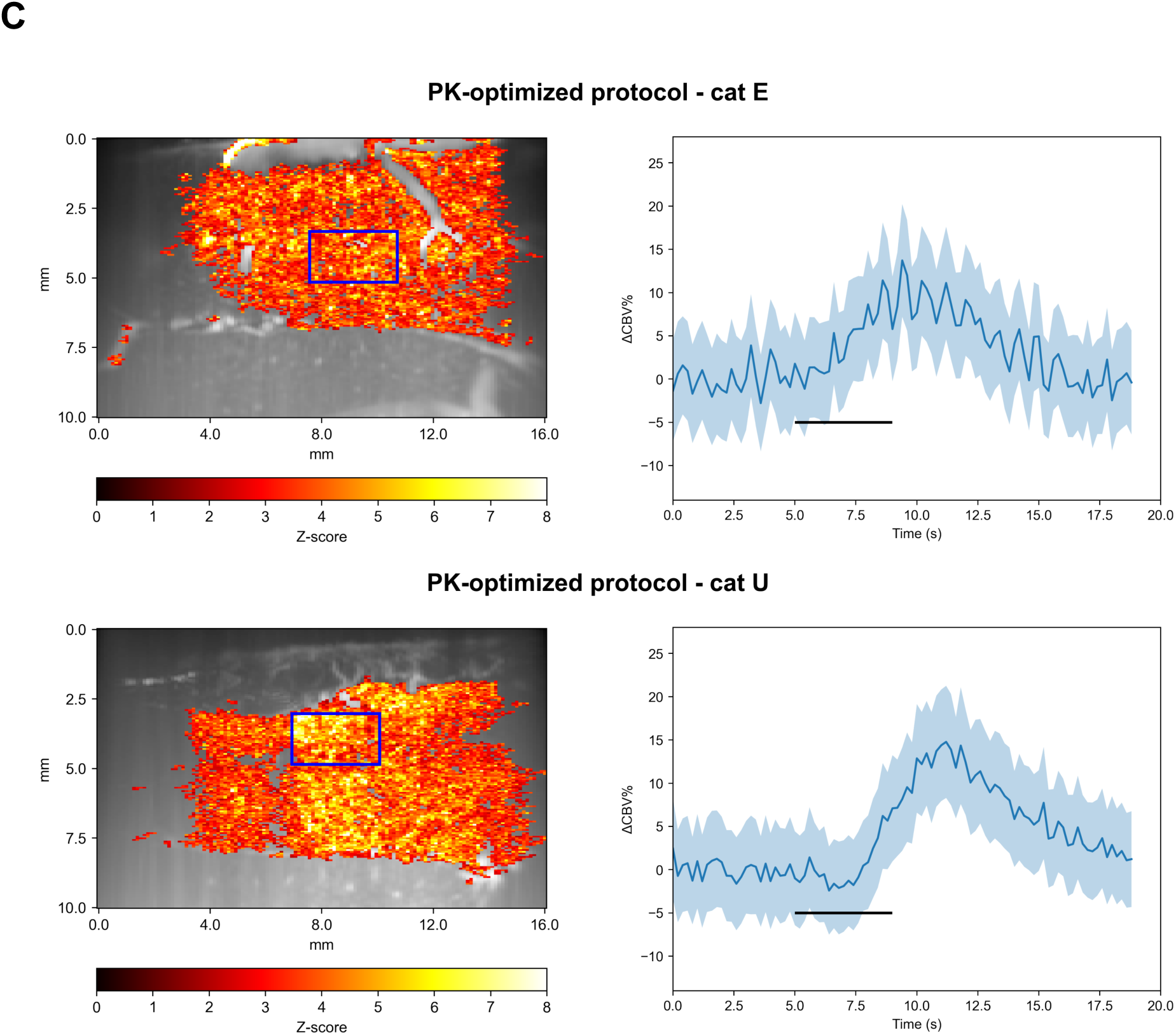
Optimizing the anesthesia protocol for medetomidine-sensitive individual animals. **(A)** Representative response maps of a normal (top, cat U) and a medetomidine-sensitive (bottom, cat E) animal, elicited by a full-screen flickering checkerboard stimulus. The imaging plane was aimed to be the same (right hemisphere, appr. 2 mm from the midline) in both cats but individual differences in the vascular map are clearly seen in the ultrasound images. **(B)** Pharmacokinetic (PK, plasma concentration) modeling of medetomidine effect using the Brown et al. 2014 (blue) and the PK-optimized (orange) anesthesia protocol. **(C)** Representative response map of the same normal animal (cat U) as in **(A)** but with the PK- optimized anesthesia protocol, elicited by a full-screen flickering checkerboard stimulus. The imaging plane was aimed to be the same for **(C)** as in **(A),** but **(A)** and **(C)** were recorded on different days several months apart. Thus, the individual vascular maps could have changed as seen in the ultrasound images.

For the within-session and across-session stability analyses (Figure 3), visual stimulation consisted of a full-screen drifting square-wave grating presented in vertical orientation. The spatial frequency of the grating was 0.125 cycles/visual degree, drifting perpendicularly to its orientation at a temporal frequency of 2 Hz, with the direction of motion reversing every 0.5 s. The trial began with a 16 s uniform gray screen, followed by 8 s of grating presentation. For the analyses presented in Figure 3, the stimulus was presented as a single, non-repeated trial, allowing quantification of the initial onset response to visual stimulation.

**Figure 3.**
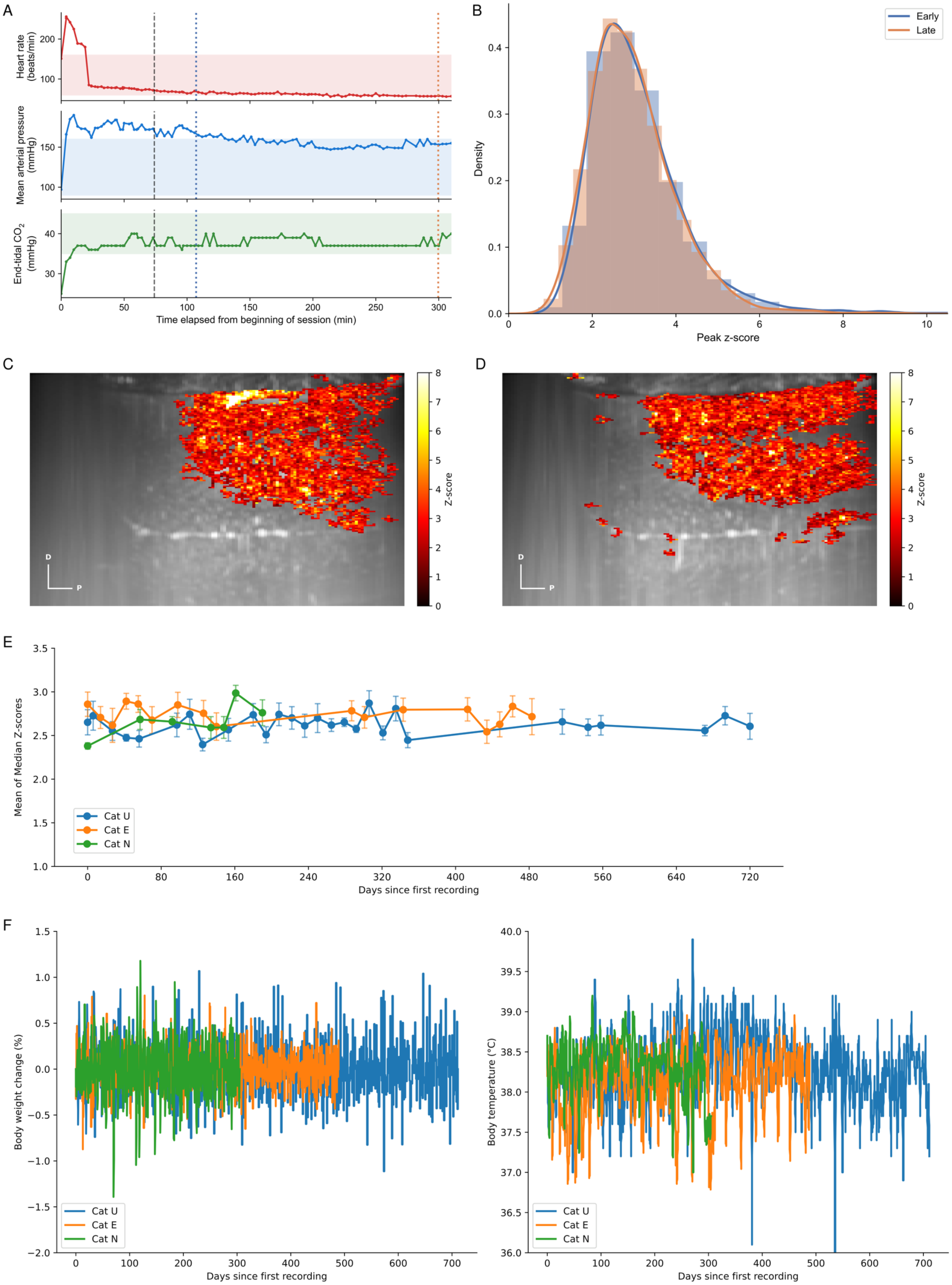
Anesthesia protocol results in physiological and functional stability throughout an experimental session and across months of recordings. (A) Evolution of physiological parameters in a single experimental session during 6 hours from the start of anesthesia monitoring. Parameters shown from top to bottom: heart rate (beats/min), mean arterial pressure (mmHg), and end-tidal CO₂ (mmHg). Vertical dashed line marks the criterion for when the functional recordings can be started (see Methods for more details). The transient period before this point reflects the induction phase of anesthesia. The two dotted vertical lines indicate the time points of the early and late recordings shown in (B– D). (B) Comparison of fUSI signal strength between two recordings made 3 hours apart from the session shown in (A). Histograms show the distribution of peak z-scores across all significantly activated pixels for an early (blue) and late (orange) recording obtained within the same session using an identical full-screen visual stimulation protocol. Solid curves show kernel density estimates (KDE) for each distribution. Distributions were statistically equivalent (two one-sided tests, TOST; equivalence margin ±0.3 z-score units; p = 4.36 × 10-11; Cohen’s d = 0.15). (C) Peak z-score activation map of the early recording from (B), overlaid on the anatomical image. Pixels not showing significant stimulus-evoked activation are masked (see Methods). (D) Same as (C) for the late recording from (B). (E) Stability of visual onset response strength across years of recordings in three animals. Each data point represents the within-day mean of the medians of peak z-score values across all significantly activated pixels from a single recording session, plotted as a function of days elapsed since first recording. Error bars denote ±1 SD. (F) Stability of body weight (left) and body temperature (right) of the n = 3 animals included in the response stability analysis. The direction indicators in (C) and (D) also serve as scale bars; each arm represents 1 mm. D: dorsal, P: posterior, TOST: two one-sided tests

### Calculating the number of significantly responding pixels

In each functional ultrasound image pixel, we compared signals along the response time course in a time window (5 sec) before stimulus onset and, to account for the hemodynamic delay in fUSI, a time window (5 sec) beginning 3 sec after stimulus onset using a one-sided Mann-Whitney U test and obtained the p-value of the test for each pixel. In the analyses for Figures 1-2 and 4, a pixel was considered significant if p < 3.2 x 10-6 which corresponds to p < 0.05 after Holm-Bonferroni correction for multiple comparisons (Holm 1979). For the analysis in Figure 3, a pixel was considered significant if p < 0.05. In this analysis, no correction for multiple comparisons was applied; rather than imposing a stringent inclusion criterion that risks excluding genuinely responsive pixels, this threshold was deliberately chosen to define a broad pixel population, such that any contamination by inactive pixels would be distributed equivalently across sessions and would therefore not confound the comparison of peak z-score distributions used to assess the stability of the onset response. For all figures, significantly responding pixels were colored according to their computed z-score in the resulting z-score map and the number of these pixels was used for comparing the signal quality in the different conditions.

### Computing average CBV response curves

Average cerebral blood volume (CBV) response curves were computed by selecting a 30-by-30-pixel region of interest (ROI) on the significant activation map in a cortical area that showed significantly responding pixels (Figure 1A left column, blue rectangle on the activation maps). Within this ROI, the pixel values were averaged and plotted along the time course of the baseline before, stimulation and baseline after periods that were defined above. For Figure 4, the ROI was selected on the activation map of the awake condition, and the exact same ROI was then used to compute similar average CBV response curves for the awake and the isoflurane-ketamine-medetomidine anesthesia conditions (Figure 4A). For Figure 4, the awake map was used as reference for ROI selection because it provides the most reliable activation pattern, and this approach avoids circular selection within the anesthesia condition.

**Figure 4.**
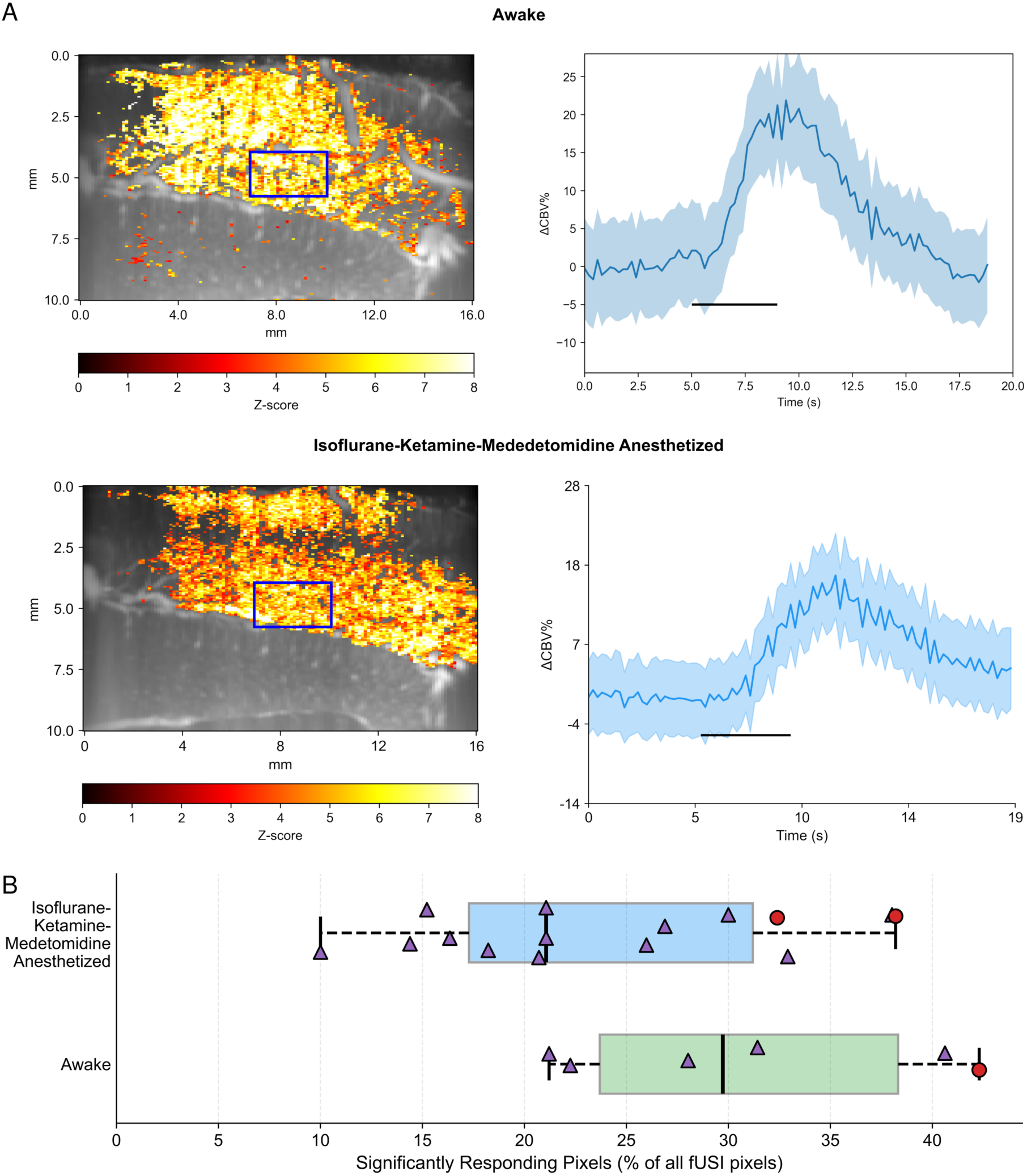
Isoflurane-ketamine-medetomidine anesthesia protocol was compatible with recording of visually evoked responses in the awake cat with functional ultrasound imaging. **(A)** Representative awake (top) and anesthetized (bottom) response maps and average hemodynamic response time courses from a cat elicited by a full-screen flickering checkerboard stimulus. **(B)** Number of significantly responding pixels as percent of all pixels (166*128=21,248) in anesthetized and awake conditions. Boxes show median and interquartile range (IQR, 25th– 75th percentile); whiskers extend to the most extreme value within 1.5×IQR of the box edges; outliers beyond this range are not displayed. Individual points show single recordings, jittered vertically and coded by experimental animal; n = 6 recordings/3 days awake, n = 15 recordings/14 days anesthetized, two animals). Due to the low number of recordings, especially in the awake condition, no formal statistical test was carried out.

### Comparing the number of significantly responding pixels across anesthesia conditions

In Figures 1B and 4B, the number of significantly responding pixels – expressed as percent of all pixels in a fUS image (166*128=21,248 pixels) – is represented on a descriptive boxplot where boxes show median and interquartile range (IQR, 25th–75th percentile); whiskers extend to the most extreme value within 1.5×IQR of the box edges; and outliers beyond this range are not displayed. Individual points show single recordings, jittered vertically and coded by experimental animal. No formal statistical analysis was carried out due to the low number of recordings in the alfaxalone-ketamine-medetomidine and isoflurane-ketamine-fentanyl conditions.

### Pharmacokinetic modeling

To assess the effect of medetomidine in the sensitive cat, and since we had no access to medetomidine plasma concentration measurements in the lab, we implemented a pharmacokinetic model for medetomidine plasma concentration, based on medetomidine pharmacokinetic parameters published for cats. A one-compartment open model was used to describe medetomidine plasma pharmacokinetics. The absorption from intramuscular (IM) injection sites was modeled as a first-order process with absorption rate constant Ka = 0.075 min⁻¹ (corresponding to a plasma Tmax of approximately 30 min, consistent with reported feline IM kinetics). Systemic clearance and volume of distribution were taken from published feline PK data (CL = 13.3 mL/min/kg; Vd = 2.98 L/kg) (Salonen 1989; Pypendop et al. 2011; Pypendop and Ilkiw 2014; Fernandes et al. 2024), with the elimination rate constant k_el_ derived as k_el_ = CL/Vd. Except for the (Salonen 1989) paper, all other publications report values for dexmedetomidine instead of medetomidine. Dexmedetomidine is the pharmacologically active enantiomer of medetomidine, so published dexmedetomidine values were adjusted for medetomidine based on the common double-volume dosing of medetomidine compared to dexmedetomidine to get the same effect. For IV bolus or IV CRI administration, absorption-site kinetics were omitted, and drug was assumed to enter the central compartment directly. The ordinary differential equation system constructed with the given parameters was integrated numerically using a 4th/5th-order Runge–Kutta solver (RK45; SciPy solve_ivp) at 1-minute resolution.

To optimize the anesthesia protocol, a step-wise full-IV CRI schedule was designed by simulating the full PK time course from the initial bolus through the planned experiment duration (Figure 2B, orange, PK-optimized protocol). CRI rates were adjusted over multiple experimental sessions to achieve adequate sedation without excessive suppression of hemodynamic responses.

### PK-optimized isoflurane-ketamine-medetomidine anesthesia

On the day before the anesthetized experiment, an IV cannula was placed in one of the cephalic or saphenous veins of the cat in light sedation (0.25 mg/kg midazolam + 20 µg/kg medetomidine injected IM). On the experiment day, the cat was premedicated with 0.25 mg/kg midazolam (Dormicum, Egis), 1 mg/kg alfaxalone (Alfaxan Multidose, Jurox) and 0.5 mg/kg ketamine (CP-Ketamine, Produlab Pharma) injected intravenously. Further alfaxalone was given intravenously (0.5-1.2 mg/kg) until the corneal and palpebral reflexes were absent. After intubation, the cat was put in a custom-built head holder that connects to the 3D-printed skull implant and enables pain-free head fixation. Medetomidine was induced with a 5 µg/kg bolus given intravenously in 5 minutes. Anesthesia was maintained with 1% isoflurane, 0.6 mg/kg/h ketamine and 3 µg/kg/h medetomidine infusion during the installation of the functional ultrasound transducer. Isoflurane rate was reduced to 0.5% in 0.1% steps every 5 minutes, while ketamine rate was increased to 0.72-0.84 mg/kg/h and medetomidine rate was reduced to 2 µg/kg/h, both in a single step, for fUSI. The first fUSI recording was performed at least 10 minutes after reaching the 0.5% isoflurane rate for imaging.

### Direct comparison of response strength to evaluate functional ultrasound response stability in the optimal isoflurane-ketamine-medetomidine anesthesia

To assess the stability of the onset response across the recording session, we compared the distribution of per-pixel peak z-scores between an early and a late recording acquired on the same day. For each recording, the onset response was quantified for all significantly activated pixels. For each significantly activated pixel, the hemodynamic signal was z-scored using a baseline standard deviation computed from the baseline frames, and the baseline mean was subtracted prior to z-scoring. The peak z-score within a 5 s window following stimulus onset was then extracted per pixel. To test whether the two distributions were equivalent rather than merely non-significantly different, we applied a two one-sided t-test procedure (TOST; Welch’s approximation for unequal variances) with an equivalence margin of ±0.3 z-score units. Cohen’s d was calculated as a measure of effect size, using the pooled standard deviation of the two samples.

### Longitudinal stability of onset response strength in the optimal isoflurane-ketamine-medetomidine anesthesia

To assess the stability of the visual onset response over months to years of recordings, the median peak z-score across all significantly activated pixels was computed for each eligible recording and the daily means of median z-scores were plotted as a function of days elapsed since first recording. All anesthetized recordings from the two subject animals that used the identical full-screen visual stimulation protocol were included from every session with at least two such recordings. For each experimental day the within-day mean and standard deviation of the median z-scores were computed. The resulting time series was plotted separately for each animal, with individual recording values shown alongside session summary statistics, to visualize the trajectory of response strength across the full longitudinal dataset.

### Stability of body weight and body temperature of the experimental animals

The body weight and rectal body temperature of the n = 3 animals included in the onset response stability analysis was measured once every day, including the anesthetized experiment days when these were measured in the awake animal before premedication for anesthesia. Body weight (left) and temperature (right) values are plotted in Figure 3F, using a 7-day moving average for visual smoothing. To assess whether repeated anesthetized procedures had cumulative effects on animal health, we fitted linear mixed-effects models (LME) using restricted maximum likelihood (REML) estimation (Python statsmodels v0.14.6). Body weight change (% from previous recording) and body temperature (°C, measured from awake animals prior to anesthesia) were modeled as outcome variables in separate models. Days since first recording served as the fixed effect of interest in time-trend models. Anesthetized sessions were identified automatically from procedure logs as entries containing “anesthetized” in the comment field (total: 51 sessions across 3 animals). A binary anesthesia flag (1 = within ±5 days of a session date, 0 = otherwise) served as the fixed effect in anesthesia-effect models. All models included a random intercept per animal to account for repeated measures. For the anesthesia-effect model on body temperature, an additional random intercept per individual session was included to account for between-session variability in baseline temperature unrelated to anesthesia. Statistical significance was assessed at α = 0.05.

## Results

### Comparing anesthesia protocols for functional ultrasound imaging

We recorded visually evoked functional ultrasound responses from the visual cortex of three cats, in different anesthesia protocols. Isoflurane-ketamine-medetomidine anesthesia was chosen to be the reference because it has been published to provide good functional neuroimaging responses in cats (Brown et al. 2014) and marmosets (Ortiz-Rios et al. 2024) and because it also provided good functional responses in our recordings. Alfaxalone-ketamine-medetomidine and isoflurane-ketamine-fentanyl anesthesia have also been published to provide functional neuroimaging responses in cats (Levine et al. 2020) and macaques (Logothetis et al. 2001), respectively, so we chose these two methods to compare to the isoflurane-ketamine-medetomidine anesthesia. Alfaxalone-ketamine-medetomidine recordings yielded a lower number of significantly responding pixels than during isoflurane-ketamine-medetomidine anesthesia (median = 0.9% vs. median = 21.1%). Isoflurane-ketamine-fentanyl recordings also yielded a lower number of significantly responding pixels than during isoflurane-ketamine-medetomidine anesthesia (median = 4.4% vs. median = 21.1%). Due to the low number of recordings in the alfaxalone and fentanyl conditions, no formal statistical test was carried out. The wide confidence intervals reflect the limited number of recordings in the alfaxalone-ketamine-medetomidine and isoflurane-ketamine-fentanyl conditions. In one isoflurane-ketamine-fentanyl recording, isoflurane was reduced to 0.3%. This provided the highest number of significantly responding pixels for this protocol (see the highest, outlier value marked by an asterisk on panel B, Isoflurane-Ketamine-Fentanyl). However, reducing isoflurane to 0.3% quickly led to an unstable anesthesia reflected by a rapidly increasing heart rate and mean arterial pressure. Taken together, our recordings indicate that neither alfaxalone-ketamine-medetomidine nor isoflurane-ketamine-fentanyl anesthesia consistently provide robust visually evoked responses under the tested conditions while providing a stable anesthesia at the same time.

### Optimizing isoflurane-ketamine-medetomidine anesthesia for individual medetomidine sensitivity

Medetomidine is a potent vasoconstrictor and it is well-documented in cats that the single IM dose we used (20 µg/kg) can already induce severe bradycardia (heart rate < 100 bpm) in sensitive cats (Lamont et al. 2001; Pypendop et al. 2011). This individual difference in medetomidine sensitivity is reflected in the lack of visually-evoked functional ultrasound response (Figure 2A) in a medetomidine-sensitive cat (as indicated by its more severe, < 70 bpm heart rate after a single 20 µg/kg IM medetomidine injection; heart rate data not shown). To assess the effect of medetomidine in the medetomidine-sensitive cat, we implemented a pharmacokinetic model of medetomidine plasma concentration. The pharmacokinetic parameters (Vd, CL) were drawn from the published literature (Salonen 1989; Pypendop et al. 2011; Pypendop and Ilkiw 2014; Fernandes et al. 2024) due to the lack of medetomidine plasma samples from our experimental animals. Using the pharmacokinetic model and the experimental observations of anesthesia stability and functional ultrasound response quality as guides, an optimal anesthesia protocol including full-IV medetomidine dosing was found that provided a stable anesthesia and functional ultrasound signal at the same time (Figure 2C, top). The stepwise isoflurane dose reduction, as well as the careful IV medetomidine dosing in this optimal protocol ensures that cardiovascular stability (heart rate > 75 bpm; data not shown) is maintained that also contributes to the functional ultrasound signal quality (Figure 2C, top). The optimal anesthesia protocol generalizes to the nomral cat as well (Figure 2C, bottom); thus, a single optimal anesthesia protocol could be used for all cats involved in the study. The stability of the PK-optimized anesthesia and functional ultrasound signals are further evaluated in Figure 3.

### Physiological and functional stability of the PK-optimized isoflurane-ketamine-medetomidine anesthesia protocol throughout an experimental session and across months of recordings

A central requirement for longitudinal fUSI studies is that the anesthesia protocol maintains stable physiology and does not introduce systematic drift in hemodynamic signal strength across a recording session or over the lifetime of the implant. We therefore evaluated both within-session and long-term stability of our preparation.

To directly test whether fUSI signal strength was preserved across a recording session, we compared the onset responses of two recordings acquired 3 hours and 12 minutes apart within the same session, using an identical full-screen drifting grating stimulus presented once per recording. For each recording, the peak z-score within the first 5 s of the stimulus onset response was computed for every significantly activated pixel, and the resulting distributions were compared using equivalence testing (TOST). The distributions of peak z-scores were statistically equivalent between the early and late recordings (TOST, equivalence margin ±0.3 z-score units; p = 4.36 × 10⁻¹¹; Cohen’s d = 0.15; Figure 3B), with the small effect size confirming that any difference between sessions was negligible in magnitude.

To evaluate stability over a longer timescale, we tracked the median peak z-score of the onset response across all eligible anesthetized recording sessions in three animals, spanning 23 months for cat U, 16 months for cat E and 6 months for cat N from the first functional recording (Figure 3E). Across a total of 464 recordings (cat U: 262, cat E: 155, cat N: 47) the median onset response remained stable without systematic increase or decrease over time in either animal. A linear mixed-effects model with days since first recording as the fixed effect of interest and random intercepts for animal and recording session confirmed the absence of a significant temporal trend in median onset response strength across the full longitudinal dataset (slope = 9 × 10⁻6 z-score units/day, 95% CI [−1.8 × 10⁻⁴, 1.9 × 10⁻⁴]; p = 0.928; N = 464 recordings, 51 sessions, 3 animals). Together, these results demonstrate that the anesthesia protocol supports both within-session and long-term stability of fUSI signal quality, validating its use for longitudinal experiments.

### Stability of body weight and body temperature of the experimental animals

Body weight and body temperature were monitored across up to two years of bi-weekly anesthetized procedures (N = 3 cats, total N = 1,620 daily observations). All values remained within normal feline physiological ranges throughout the study period (Figure 3F; no animal required veterinary intervention for weight loss, fever, or hypothermia attributable to the protocol).

#### Time trends

Body weight change showed no significant trend over time (β = 0.0000%/day, SE < 0.001, p = 0.936), indicating stable body weight across the full monitoring period. Body temperature likewise showed no significant trend over time (β = 0.0000°C/day, SE < 0.0001, p = 0.773), indicating no cumulative drift in resting body temperature across up to two years of repeated procedures.

#### Effect of repeated anesthetized procedures

Body weight change was not significantly associated with anesthetized procedure days (β = −0.075%, SE = 0.096, p = 0.432), and this result was unchanged when an additional random intercept for individual anesthesia session was included (β = −0.073%, SE = 0.094, p = 0.438; session variance component ≈ 0, indicating no meaningful between-session heterogeneity in weight). Body temperature, including pre-induction body temperature – measured from awake animals before each session and therefore unaffected by intra-procedural temperature management – was also not significantly associated with anesthetized procedures (β = −0.032°C, SE = 0.023, p = 0.162). When an additional random intercept for individual session was included, the effect remained non-significant (β = −0.028°C, SE = 0.137, p = 0.839), with a modest between-session variance component (σ²_session = 0.053, SE = 0.041) reflecting ordinary day-to-day variation in resting temperature unrelated to anesthesia history. Together, these results indicate that repeated bi-weekly anesthesia over up to two years did not produce detectable cumulative or acute effects on body weight or resting body temperature in any of the three animals.

### Comparing the isoflurane-ketamine-medetomidine anesthesia to wakefulness

As an initial, exploratory characterization of how anesthetized fUSI responses relate to the awake state, we recorded visually evoked functional ultrasound responses from the visual cortex of two cats, in wakefulness and anesthesia. Both awake and anesthetized conditions yielded clear visually evoked responses, as illustrated by the representative z-score maps and CBV time courses in Figure 4A. Due to the low number of recordings, especially in the awake condition, no formal statistical test was carried out. The boxplots (Figure 4B) clearly indicate a reduction in the number of significantly responding pixels in anesthesia (median = 21.1%) compared to the awake condition (median = 29.7%). However, along with the preserved activation map and response curve, this result indicates that the presented anesthesia protocol enables the recording of visually evoked fUSI responses that are comparable to responses recorded in the awake state.

## Discussion

The classical functional neuroimaging method, fMRI, mostly relies on anesthesia in large-brained animal models. There have been multiple anesthesia protocols published for fMRI in large animals. Many of these use the combination of isoflurane and one or more injectable anesthetics, such as ketamine and medetomidine (Brown et al. 2014) or fentanyl (Logothetis et al. 2001) but a purely injectable-anesthetic-based protocol has also been published for cats (Levine et al. 2020). When performing fUSI on large-brained animals, these protocols can be used as candidates, but since fMRI and fUSI rely on different aspects of the same neurovascular-coupling-driven signal (BOLD for fMRI vs CBV for fUSI), it is not guaranteed that any of these protocols will readily translate for fUSI as well. In this study, we present our non-extensive screening of some of the published fMRI anesthesia protocols evaluated for fUSI (Figure 1), and we show that the isoflurane-ketamine-medetomidine protocol provides the most reliable fUSI responses among the screened. This aligns well with the results of (Levine et al. 2020) where they showed that a similar anesthesia protocol provided higher amplitude fMRI responses in the cat visual and auditory cortices than an alfaxalone-based protocol. The isoflurane-ketamine-fentanyl protocol, while achieving higher pixel counts at reduced isoflurane concentrations, produced hemodynamic instability that precluded its reliable use, consistent with the known narrow therapeutic window of fentanyl in cats (Bortolami and Love 2015).

Medetomidine is a potent vasoconstrictor and central sympathetic inhibitor. As such, it has already been shown that domestic cats can have highly variable cardiovascular responses to medetomidine, including severe bradycardia (heart rate < 100 bpm) (Lamont et al. 2001; Pypendop et al. 2011). Excessive bradycardia is highly undesirable for fUSI due to its cerebral-perfusion-reduction effect that can lead to the loss of fUSI signal. We observed such a fUSI signal loss in one of the cats involved in this study. Domestic cats are an outbred species with substantial inter-individual genetic variation (Anderson et al. 2022; Lipinski et al. 2008; Montague et al. 2014), and our colony, bred in-house and sourced from licensed suppliers in accordance with EU and Hungarian regulations, reflects this natural genetic diversity. As demonstrated by (Ding et al. 2023) in humans and for dexmedetomidine (the active enantiomer in racemic medetomidine), the hemodynamic response to medetomidine has a strong genetic background, which likely explains the inter-individual differences in medetomidine sensitivity we observed. To mitigate this effect, we used a PK model based on published feline parameters (Fernandes et al. 2024; Pypendop and Ilkiw 2014; Salonen 1989) to guide development of a medetomidine-infusion regimen, with physiological stability and functional ultrasound signal quality used as experimental readouts (Figure 2). Using this approach, we optimized the isoflurane-ketamine-medetomidine anesthesia protocol by replacing a single medetomidine intramuscular bolus with a low-rate medetomidine infusion. This way, there is no medetomidine plasma-concentration and effective-drive peak after a single bolus (Figure 2B) that presumably leads to the suppressive effects of medetomidine in sensitive animals. The beneficial effect of the medetomidine infusion is reflected by the preserved functional ultrasound signal in the sensitive animal (Figure 2C). The PK-optimized protocol also generalizes to the normal animal (Figure 2C), providing an optimal anesthesia protocol for visual cortical fUSI in cats. This medetomidine-infusion-based optimal protocol also aligns well with the recently published anesthesia protocol for fMRI in marmosets (Ortiz-Rios et al. 2024), indicating that our protocol could be adopted for other large-brained species as well.

We demonstrate that the optimal anesthesia protocol permits multiple-hour-long fUSI experiments that are repeatable for up to 23 months in a single animal (Figure 3). Long-term stability of the animal preparation is also enabled by our custom-designed skull chamber preparation that will be described in detail in a separate publication. Longitudinal experiments in a single animal carry multiple benefits; they significantly reduce the number of animals used for research, while enabling the study of how brain structure and function changes with time, for example through learning and experience. The reproducibility of fUSI signals that is enabled by our PK-optimized anesthesia protocol can facilitate the study of such long-term brain structure and function changes that can happen between anesthetized recording sessions.

A substantial advantage of fUSI over fMRI is that fUSI enables awake recordings in large-brained animals (Blaize et al. 2020), whereas awake fMRI, while not impossible, it still limited in such species (Basso et al. 2021; Goense et al. 2007; Szabó et al. 2019). Motivated by this feature of fUSI, we compared the fUSI signals recorded with our PK-optimized anesthesia protocol to awake recordings in the same animals. The qualitative similarity between anesthetized and awake fUSI responses (Figure 4) is encouraging, although formal equivalence could not be statistically demonstrated (TOST p = 0.472), most likely due to insufficient statistical power from the low number of awake recording days in untrained animals rather than a true difference between conditions. The observed pixel counts are numerically consistent with preserved anesthetized responses (awake: 6,581 ± 1,904 pixels; anesthetized: 5,120 ± 1,867 pixels, Figure 4B). A definitive equivalence study would require a larger cohort of animals trained for awake head-fixed recordings – a goal for future work.

### Long-term physiological monitoring

“Longitudinal anesthetized imaging may provide a complementary approach when experiments require multi-hour acquisition, controlled visual stimulation, or gaze stabilization. To examine whether repeated procedures were associated with changes in readily monitored physiological measures, we tracked body weight and pre-induction body temperature throughout the study. Neither measure showed a significant temporal trend or a significant association with anesthetized procedure days across the three animals. Together with the stable physiological parameters observed during multi-hour recordings, these data support the physiological tolerability of the present preparation over the monitored period.

The protocol also maintained visual response strength across repeated recording sessions, with no detectable temporal trend over the longitudinal dataset. Thus, within the limits of the small cohort and the physiological measures assessed, the preparation supports repeated functional ultrasound imaging over months in the cat visual cortex. This feature is particularly valuable for experiments requiring longitudinal or large-volume functional mapping that would be difficult to complete within shorter awake recording sessions.

Our study is limited to a low number of animals (n=3), however, it still indicates the potential of long-term reproducible fUSI recordings in the visual cortex of cats. While we only present data from the visual cortex of cats, studies from other labs (Brown et al. 2014; Levine et al. 2020) that used a similar isoflurane-ketamine-medetomidine-based anesthesia as discussed above suggest that our PK-optimized anesthesia protocol could also be applied for imaging from other cat cortical areas, such as the auditory cortex. Functional mapping of the cerebral cortex using neuroimaging in large-brained animal species opens the way for a deeper understanding of complex cortical structures. Functional ultrasound imaging has already been applied to reconstruct large-volume 3D functional maps in the awake ferret auditory cortex (Bimbard et al. 2018) and also has been demonstrated to provide large-scale functional maps in the awake macaque visual cortex (Blaize et al. 2020). However, as brain volume increases with increasing brain complexity, the reconstruction of large-volume cortical functional maps with fUSI requires increasingly more time due to the relatively low temporal frequency (< 10 Hz) of fUSI, while the awake preparations are limited in acquisition time to promote animal welfare. In this context, a stable and reproducible anesthesia protocol that we present in this study can enable the acquisition of large-volume functional maps in large-brained species, such as cats and macaques. Moreover, our chronic skull implant and reproducible anesthesia protocol might enable simultaneous fUSI and electrophysiological recordings. Although simultaneous fUSI and single unit electrophysiological recordings have already been published in awake macaques, (Claron et al. 2023), they might be limited by the constraints of awake, behavioral recordings. For example, the spatial coverage of simultaneous fUSI and electrophysiology was limited to what could be acquired within the 1-hour awake behavioral window in the (Claron et al. 2023) study. Our anesthesia protocol, by removing this time constraint, opens the possibility of extending such experiments to substantially larger cortical volumes, enabling a more complete mapping of the neural-hemodynamic relationship across the cerebral cortex.

## Acknowledgements

The authors gratefully acknowledge Prof. Botond Roska (IOB, Basel) for making the functional ultrasound recording setup available for this study as well as Alan Urban (VIB, Leuven) for advice. The authors would like to thank Sarolt Kinga Gintner, Renáta Radics, Luca Benedek, Zsombor Fülei and Hanna Orvos-Nagy for their assistance in animal care. The authors also would like to thank Beatrix Kovács, Fanni Somogyi for their extensive support in performing surgeries and experiments, and for Gábor Ábrahám, Hella Czigány, Virág György, Dávid Jólesz, Virág Sári and Fanni Soós for their assistance during the experiments.

This work has been supported by 2019-2.1.7-ERA-NET-2021-00047, the Lendület (“Momentum”) Programme of the Hungarian Academy of Sciences, Excellence 151368 from Ministry of Innovation and Technology of Hungary (NRDI fund), CELSA/24/020 and KSZF-161/2024 to DaH, 2024-2.1.2-EKÖP-KDP New National Excellence Program of the Ministry for Culture and Innovation (NKFIH) to KCs.

## Notes

### Competing Interest Statement

The authors have declared no competing interest.

## References

Anderson, Heidi, Stephen Davison, Katherine M. Lytle, et al. 2022. “Genetic Epidemiology of Blood Type, Disease and Trait Variants, and Genome-Wide Genetic Diversity in over 11,000 Domestic Cats.” PLoS Genetics 18 (6): e1009804. 10.1371/journal.pgen.1009804.

Basso, M. A., S. Frey, K. A. Guerriero, et al. 2021. “Using Non-Invasive Neuroimaging to Enhance the Care, Well-Being and Experimental Outcomes of Laboratory Non-Human Primates (Monkeys).” NeuroImage 228 (March): 117667. 10.1016/j.neuroimage.2020.117667.

Bimbard, Célian, Charlie Demene, Constantin Girard, et al. 2018. “Multi-Scale Mapping along the Auditory Hierarchy Using High-Resolution Functional UltraSound in the Awake Ferret.” eLife 7 (June): e35028. 10.7554/eLife.35028.

Bini, G., K. M. Bailey, J. T. Voyvodic, L. Chiavaccini, K. R. Munana, and E. K. Keenihan. 2023. “Effects of Alfaxalone, Propofol and Isoflurane on Cerebral Blood Flow and Cerebrovascular Reactivity to Carbon Dioxide in Dogs: A Pilot Study.” The Veterinary Journal 291 (January):105939. 10.1016/j.tvjl.2022.105939.

Blaize, Kévin, Fabrice Arcizet, Marc Gesnik, et al. 2020. “Functional Ultrasound Imaging of Deep Visual Cortex in Awake Nonhuman Primates.” Proceedings of the National Academy of Sciences 117 (25): 14453–63. 10.1073/pnas.1916787117.

Bortolami, Elisa, and Emma J. Love. 2015. “Practical Use of Opioids in Cats: A State-of-the-Art, Evidence-Based Review.” Journal of Feline Medicine and Surgery 17 (4): 283–311. 10.1177/1098612X15572970.

Brown, Trecia A., Joseph S. Gati, Sarah M. Hughes, Pam L. Nixon, Ravi S. Menon, and Stephen G. Lomber. 2014. “Functional Imaging of Auditory Cortex in Adult Cats Using High-Field fMRI.” Journal of Visualized Experiments, no. 84 (February): 50872. 10.3791/50872.

Claron, Julien, Matthieu Provansal, Quentin Salardaine, et al. 2023. “Co-Variations of Cerebral Blood Volume and Single Neurons Discharge during Resting State and Visual Cognitive Tasks in Non-Human Primates.” Cell Reports 42 (4). 10.1016/j.celrep.2023.112369.

Ding, Yuanyuan, Aiqing Liu, Yafeng Wang, et al. 2023. “Genetic Polymorphisms Are Associated with Individual Susceptibility to Dexmedetomidine.” Frontiers in Genetics 14 (August). 10.3389/fgene.2023.1187415.

Fernandes, Naftáli S., Yanna D. B. Passos, Kathryn N. Arcoverde, et al. 2024. “Clinical Effects and Pharmacokinetic Profile of Intramuscular Dexmedetomidine (10 Μg/Kg) in Cats.” Animals : An Open Access Journal from MDPI 14 (15): 2274. 10.3390/ani14152274.

Gao, Yu-Rong, Yuncong Ma, Qingguang Zhang, Aaron T. Winder, Zhifeng Liang, Lilith Antinori, Patrick J. Drew, and Nanyin Zhang. 2016. “Time to Wake up: Studying Neurovascular Coupling and Brain-Wide Circuit Function in the Un-Anesthetized Animal.” NeuroImage 153:382. 10.1016/j.neuroimage.2016.11.069.

Goense, Jozien B. M., Anne-Catherin Zappe, and Nikos K. Logothetis. 2007. “High-Resolution fMRI of Macaque V1.” *Magnetic Resonance Imaging*, Proceedings of the International School on Magnetic Resonance and Brain Function, vol. 25 (6): 6. 10.1016/j.mri.2007.02.013.

Holm, Sture. 1979. “A Simple Sequentially Rejective Multiple Test Procedure.” Scandinavian Journal of Statistics 6 (2): 65–70.

Lamont, Leigh A., Barret J. Bulmer, Kurt A. Grimm, William J. Tranquilli, and David D. Sisson. 2001. “Cardiopulmonary Evaluation of the Use of Medetomidine Hydrochloride in Cats.” American Journal of Veterinary Research. American Journal of Veterinary Research 62 (11): 1745–62. 10.2460/ajvr.2001.62.1745.

Levine, Alexandra T., Benson Li, Paisley Barnes, Stephen G. Lomber, and Blake E. Butler. 2020. “Assessment of Anesthesia on Physiological Stability and BOLD Signal Reliability during Visual or Acoustic Stimulation in the Cat.” Journal of Neuroscience Methods 334 (March): 108603. 10.1016/j.jneumeth.2020.108603.

Lipinski, Monika J., Lutz Froenicke, Kathleen C. Baysac, et al. 2008. “The Ascent of Cat Breeds: Genetic Evaluations of Breeds and Worldwide Random-Bred Populations.” Genomics 91 (1): 12–21. 10.1016/j.ygeno.2007.10.009.

Logothetis, Nikos K., Jon Pauls, Mark Augath, Torsten Trinath, and Axel Oeltermann. 2001. “Neurophysiological Investigation of the Basis of the fMRI Signal.” Nature 412 (6843): 150–57. 10.1038/35084005.

Macé, Emilie, Gabriel Montaldo, Ivan Cohen, Michel Baulac, Mathias Fink, and Mickael Tanter. 2011. “Functional Ultrasound Imaging of the Brain.” Nature Methods 8 (8): 662–64. 10.1038/nmeth.1641.

Montague, Michael J., Gang Li, Barbara Gandolfi, et al. 2014. “Comparative Analysis of the Domestic Cat Genome Reveals Genetic Signatures Underlying Feline Biology and Domestication.” Biological Sciences. Proceedings of the National Academy of Sciences 111 (48): 48. 10.1073/pnas.1410083111.

Montaldo, Gabriel, Alan Urban, and Emilie Macé. 2022. “Functional Ultrasound Neuroimaging.” Annual Review of Neuroscience 45 (1): 491–513. 10.1146/annurev-neuro-111020-100706.

Ogawa, S., T. M. Lee, A. R. Kay, and D. W. Tank. 1990. “Brain Magnetic Resonance Imaging with Contrast Dependent on Blood Oxygenation.” Proceedings of the National Academy of Sciences of the United States of America 87 (24): 9868–72. 10.1073/pnas.87.24.9868.

Ortiz-Rios, Michael, Nikoloz Sirmpilatze, Jessica König, and Susann Boreitus. 2024. “An Anesthetic Protocol for Preserving Functional Network Structure in the Marmoset Monkey Brain.” Imaging Neuroscience 2 (July): imag–2–00230. 10.1162/imag_a_00230.

Peirce, Jonathan W. 2007. “PsychoPy—Psychophysics Software in Python.” Journal of Neuroscience Methods 162 (1): 8–13. 10.1016/j.jneumeth.2006.11.017.

Porchet, H. C., N. L. Benowitz, and L. B. Sheiner. 1988. “Pharmacodynamic Model of Tolerance: Application to Nicotine.” The Journal of Pharmacology and Experimental Therapeutics 244 (1): 231–36.

Pypendop, Bruno H., Linda S. Barter, Scott D. Stanley, and Jan E. Ilkiw. 2011. “Hemodynamic Effects of Dexmedetomidine in Isoflurane-Anesthetized Cats.” Veterinary Anaesthesia and Analgesia 38 (6): 555–67. 10.1111/j.1467-2995.2011.00663.x.

Pypendop, Bruno H., and Jan E. Ilkiw. 2014. “Pharmacokinetics of Dexmedetomidine after Intravenous Administration of a Bolus to Cats.” American Journal of Veterinary Research. American Journal of Veterinary Research 75 (5): 441–45. 10.2460/ajvr.75.5.441.

Salonen, J. S. 1989. “Pharmacokinetics of Medetomidine.” Acta Veterinaria Scandinavica. Supplementum 85: 49–54.

Sheiner, L. B., D. R. Stanski, S. Vozeh, R. D. Miller, and J. Ham. 1979. “Simultaneous Modeling of Pharmacokinetics and Pharmacodynamics: Application to d-Tubocurarine.” Clinical Pharmacology and Therapeutics 25 (3): 358–71. 10.1002/cpt1979253358.

Sumiyoshi, Akira, Robin J. Keeley, and Hanbing Lu. 2019. “Physiological Considerations of Functional Magnetic Resonance Imaging in Animal Models.” *Biological Psychiatry: Cognitive Neuroscience and Neuroimaging*, The Bridging of Scales: Techniques for Translational Neuroscience, 4 (6): 522–32. 10.1016/j.bpsc.2018.08.002.

Szabó, Dóra, Kálmán Czeibert, Ádám Kettinger, et al. 2019. “Resting-State fMRI Data of Awake Dogs (Canis Familiaris) via Group-Level Independent Component Analysis Reveal Multiple, Spatially Distributed Resting-State Networks.” Scientific Reports 9 (1): 15270. 10.1038/s41598-019-51752-2.

Urban, A., Dussaux, C., Martel, G., Brunner, C., Mace, E., Montaldo, G., 2015. Real-time imaging of brain activity in freely moving rats using functional ultrasound. Nat. Methods 12, 873–878. 10.1038/nmeth.3482

Vakkur, G. J., P. O. Bishop, and W. Kozak. 1963. “VISUAL OPTICS IN THE CAT, INCLUDING POSTERIOR NODAL DISTANCE AND RETINAL LANDMARKS.” Vision Research 3 (7–8): 289–314. 10.1016/0042-6989(63)90004-x.

Whittem, T., K. S. Pasloske, M. C. Heit, and M. G. Ranasinghe. 2008. “The Pharmacokinetics and Pharmacodynamics of Alfaxalone in Cats after Single and Multiple Intravenous Administration of Alfaxan® at Clinical and Supraclinical Doses.” Journal of Veterinary Pharmacology and Therapeutics 31 (6): 571–79. 10.1111/j.1365-2885.2008.00998.x.

